# Correcting subtle stratification in summary association statistics

**DOI:** 10.1101/076133

**Authors:** Gaurav Bhatia, Nicholas A. Furlotte, Po-Ru Loh, Xuanyao Liu, Hilary K. Finucane, Alexander Gusev, Alkes L. Price

## Abstract

Population stratification is a well-documented confounder in GWASes, and is often addressed by including principal component (PC) covariates computed from common SNPs (SNP-PCs). In our analyses of summary statistics from 36 GWASes (mean n=88k), including 20 GWASes using 23andMe data that included SNP-PC covariates, we observed a significantly inflated LD score regression (LDSC) intercept for several traits—suggesting that residual stratification remains a concern, even when SNPPC covariates are included.

Here we propose a new method, PC loading regression, to correct for stratification in summary statistics by leveraging SNP loadings for PCs computed in a large reference panel. In addition to SNP-PCs, the method can be applied to haploSNP-PCs, i.e. PCs computed from a larger number of rare haplotype variants that better capture subtle structure. Using simulations based on real genotypes from 54,000 individuals of diverse European ancestry from the Genetic Epidemiology Research on Adult Health and Aging (GERA) cohort, we show that PC loading regression effectively corrects for stratification along top PCs.

We applied PC loading regression to several traits with inflated LDSC intercepts. Correcting for the top four SNP-PCs in GERA data, we observe a significant reduction in LDSC intercept height summary statistics from the Genetic Investigation of ANthropometric Traits (GIANT) consortium, but not for 23andMe summary statistics, which already included SNP-PC covariates. However, when correcting for additional haploSNP-PCs in 23andMe GWASes, inflation in the LDSC intercept was eliminated for eye color, hair color, and skin color and substantially reduced for height (1.41 to 1.16; n=430k). Correcting for haploSNP-PCs in GIANT height summary statistics eliminated inflation in the LDSC intercept (from 1.35 to 1.00; n=250k), eliminating 27 significant association signals including one at the *LCT* locus, which is highly differentiated among European populations and widely known to produce spurious signals. Overall, our results suggest that uncorrected population stratification is a concern in GWASes of large sample size and that PC loading regression can correct for this stratification.

## Introduction

Population stratification is a well-documented confounder in genome-wide association studies (GWASes)^1-3^. While inclusion of principal component (PC) covariates computed from common single nucleotide polymorphisms (SNPs), referred to as SNPPCs, often provides an effective solution to stratification^4^, this strategy will not correct for subtle stratification that is not captured by SNP-PCs, and cannot be applied when analyzing GWAS summary statistics directly. In our analyses of summary statistics from 36 GWASes (mean sample size n=88k), including 20 GWASes using 23andMe, Inc. data that included SNP-PC covariates, we applied LD score regression^5^ to assess the existence of uncorrected stratification. We observed a significantly inflated LD score regression (LDSC) intercept for several GWASes including a number of studies that included SNPPC covariates. This suggests that population stratification, especially subtle stratification that is not captured by SNP-PCs, is a concern in analysis of GWAS summary statistics.

We propose a new method, PC loading regression, to correct for this stratification by analyzing summary statistics directly, and using SNP loadings for PCs computed in a large reference panel: the Genetic Epidemiology Research on Adult Health and Aging (GERA) data set, which we chose based on its large number of samples (n=54k) of diverse European ancestry^6^. PC loading regression produces a set of summary statistics that are not confounded by stratification along the input PCs. In addition to SNP-PCs, the method can be applied to other axes of interest including haploSNP-PCs, i.e. PCs computed from a larger number of rare haplotype variants that better capture subtle structure. In simulations based on real genotypes, we demonstrate that PC loading regression is effective in correcting for stratification along top PCs.

Applying PC loading regression to 20 sets of 23andMe GWAS summary statistics, which already included SNP-PC covariates, to correct for the top four SNP-PCs in the GERA data, we did not observe a significant reduction in LDSC intercept. However, when correcting for additional haploSNP-PCs in 23andMe GWASes, inflation in the LDSC intercept was eliminated for eye color, hair color, and skin color and substantially reduced for height (1.41 to 1.16; n=430k). Separately, when we applied our method to correct for SNP-PCs and haploSNP-PCs in height summary statistics from the Genetic Investigation of ANthropometric Traits (GIANT) consortium, the LDSC intercept was reduced from 1.35 to 1.00 (n=250k), consistent with no confounding. After this correction, 27 previously published loci were no longer genome-wide significant. Among loci that were no longer genome-wide significant, the largest reduction in association statistics occurred at the *LCT* locus, which is widely known to produce spurious signals^3^. Overall, our results suggest that PC loading regression can correct for subtle stratification in GWASes of large sample size.

## Results

### Overview of methods

GWAS summary statistics measure the strength of association between each SNP and the studied phenotype and will be inflated by uncorrected population stratification. At a single SNP, this inflation will be driven by (a) the magnitude of the population stratification (i.e. the proportion of phenotypic variance explained by a PC) and (b) the correlation between the SNP and the PC—the SNP loading. Thus, we can estimate SNP loadings for PCs in an external reference panel^7^ and regress these out of uncorrected GWAS summary statistics. The slope of this PC loading regression will give us an estimate of the magnitude of population stratification and the residuals will give us corrected statistics (see Methods for details). Notably, in addition to SNP-PCs we can correct for SNP loadings for haploSNP-PCs, i.e. PCs computed from a larger number of rare haploSNPs (haplotype variants constructed using a four-gamete test) that better capture subtle structure^8^. By regressing GWAS summary statistics against SNP-PC and haploSNP-PC loadings, we are able to correct for subtle stratification. We have released open source software that implements PC loading regression (see URLs).

### Simulations using real genotypes

#### Assessing correction effectiveness using out-of-sample PCs

To assess the validity of PC loading regression as a tool for correction for population stratification, we performed simulations using real genotypes from the GERA data set^7^, a data set of 54,000 European American individuals of diverse European ancestry^6^ (see Methods) genotyped at 608k SNPs after QC. We split this data set into two partitions (n=27k each), and computed 16 PCs—four SNP PCs and four haploSNP PCs from each of the MAF ranges [10^−4^,10^−3^), [10^−3^,0.005) and [0.005, 0.01)—in each partition. For each PC in the first partition we simulated 20 phenotypes stratified along each PC—where the PC explained 10% of phenotypic variance and there were no true genetic loci. We then computed summary statistics in the first partition without including PC covariates and applied PC loading regression using PC loadings estimated from the second partition.

PC loading regression eliminated nearly all (>90%; ≥98% in most cases) of the inflation in average LD score regression intercept along each PC, with near perfect correction for top PCs (see Table 1). Additionally, we note that our estimates of correction effectiveness are conservative, as they are based on loadings from only half of the samples, i.e. those in the second partition of the GERA dataset. Loadings from the full set of samples will be less noisy and will likely result in improved correction.

**Table 1.**
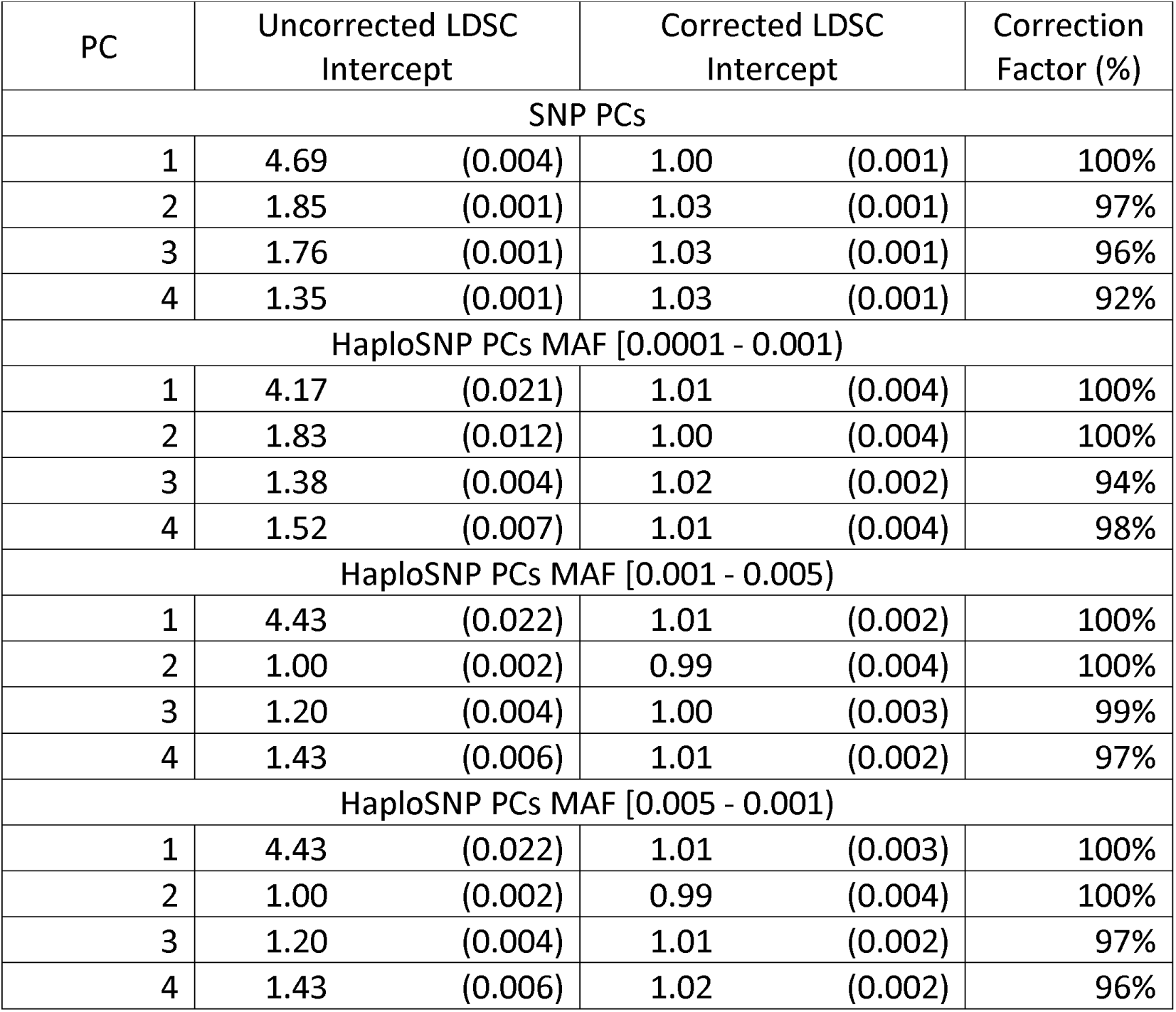
PC loading regression corrects for population stratification in simulations. We assessed the effectiveness of PC loading regression in correcting for population stratification using simulations based on the GERA dataset (n=54k). We list the mean LD score regression intercept for phenotypes stratified along each PC before and after correction with PC loading regression. This was based on correction using effect sizes for standardized genotypes (see Table S2 for per-allele corrections). We note that haploSNP PCs may not produce a large inflation in the intercept if they capture haploSNP specific structure.

| PC | Uncorrected LDSC Intercept |  | Corrected LDSC Intercept |  | Correction Factor (%) |
| --- | --- | --- | --- | --- | --- |
| SNP PCs |  |  |  |  |  |
| 1 | 4.69 | (0.004) | 1.00 | (0.001) | 100% |
| 2 | 1.85 | (0.001) | 1.03 | (0.001) | 97% |
| 3 | 1.76 | (0.001) | 1.03 | (0.001) | 96% |
| 4 | 1.35 | (0.001) | 1.03 | (0.001) | 92% |
| HaploSNP PCs MAF [0.0001 - 0.001) |  |  |  |  |  |
| 1 | 4.17 | (0.021) | 1.01 | (0.004) | 100% |
| 2 | 1.83 | (0.012) | 1.00 | (0.004) | 100% |
| 3 | 1.38 | (0.004) | 1.02 | (0.002) | 94% |
| 4 | 1.52 | (0.007) | 1.01 | (0.004) | 98% |
| HaploSNP PCs MAF [0.001 - 0.005) |  |  |  |  |  |
| 1 | 4.43 | (0.022) | 1.01 | (0.002) | 100% |
| 2 | 1.00 | (0.002) | 0.99 | (0.004) | 100% |
| 3 | 1.20 | (0.004) | 1.00 | (0.003) | 99% |
| 4 | 1.43 | (0.006) | 1.01 | (0.002) | 97% |
| HaploSNP PCs MAF [0.005 - 0.001) |  |  |  |  |  |
| 1 | 4.43 | (0.022) | 1.01 | (0.003) | 100% |
| 2 | 1.00 | (0.002) | 0.99 | (0.004) | 100% |
| 3 | 1.20 | (0.004) | 1.01 | (0.002) | 97% |
| 4 | 1.43 | (0.006) | 1.02 | (0.002) | 96% |

#### Assessing accuracy of PC loadings at imputed SNPs

PC loading regression is performed on the set of SNPs for which association statistics and PC loadings are available. To maximize this set of SNPs, we performed prephasing^9^ and imputation^10^ on the GERA reference data set using the University of Michigan Imputation Server with the Haplotype Reference Consortium^11^ as a reference panel (see Methods). To evaluate whether PC loadings were accurate at imputed SNPs, we masked 10% of genotype SNPs prior to imputation and compared PC loadings at genotyped and corresponding high-quality imputed SNPs (see Methods). Our results indicate that PC loadings at high-quality imputed SNPs are nearly identical (*r*^2^ > 0.9; *r*^2^ ≥ 0.99 in most cases) to the loadings at genotyped SNPs (see Table 2), with the exception of the lowest two PCs from the rarest haploSNP bin. Possible explanations of the discrepancy between PC loadings at genotyped and imputed SNPs are genotyping assay artifacts that are detected by lower PCs in the rarest haploSNP bin, or subtle population structure that interferes with imputation and assessment of imputation accuracy. To ensure that these issues did not impact our results, subsequent analyses only corrected for the 14 PCs with *r*^2^ > 0.9 between PC loadings from genotyped and high-quality imputed SNPs.

**Table 2.**
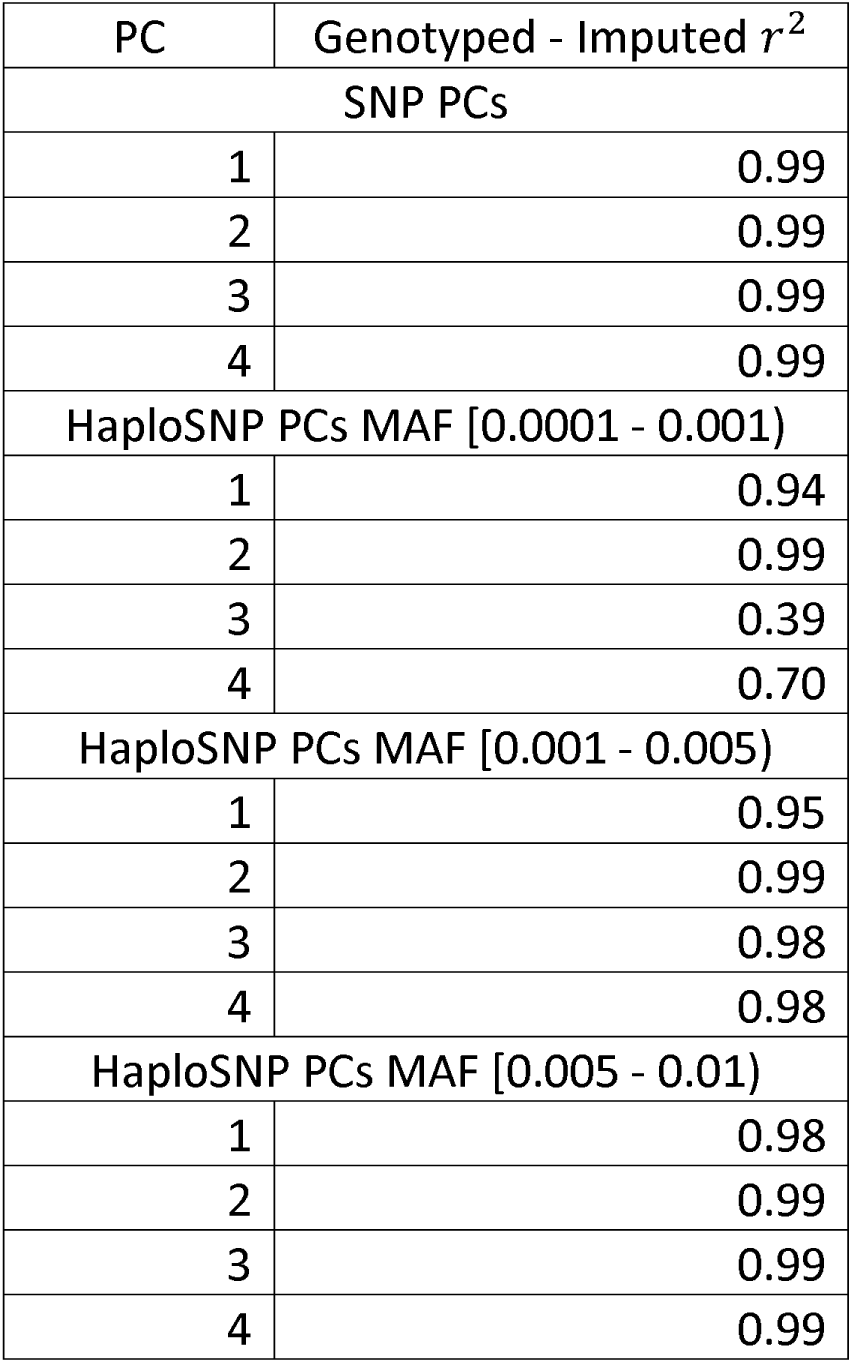
Assessing the accuracy of PC loadings at imputed SNPs. To ensure that PC loading regression could be applied at high-quality imputed SNPs, we masked 10% of genotyped SNPs from our data, performed imputation, and compared estimated PC loadings for genotyped and imputed versions of the masked SNPs. Based on this, we excluded PCs three and four of the lowest MAF bin of haploSNPs.

| PC | Genotyped - Imputed $r^2$ |
| --- | --- |
| SNP PCs |  |
| 1 | 0.99 |
| 2 | 0.99 |
| 3 | 0.99 |
| 4 | 0.99 |
| HaploSNP PCs MAF [0.0001 - 0.001) |  |
| 1 | 0.94 |
| 2 | 0.99 |
| 3 | 0.39 |
| 4 | 0.70 |
| HaploSNP PCs MAF [0.001 - 0.005) |  |
| 1 | 0.95 |
| 2 | 0.99 |
| 3 | 0.98 |
| 4 | 0.98 |
| HaploSNP PCs MAF [0.005 - 0.01) |  |
| 1 | 0.98 |
| 2 | 0.99 |
| 3 | 0.99 |
| 4 | 0.99 |

### Analysis of summary statistics from real phenotypes

#### LD Score regression intercepts of 36 GWAS summary statistics

We analyzed 36 sets of GWAS summary statistics (mean sample size n=88k), including publicly available summary statistics for 16 diseases and quantitative traits^12-24^ and summary statistics from 20 GWASes using 23andMe data that included SNP-PC covariates (see Table S1). We quantified the inflation in summary statistics due to stratification using the LD Score regression (LDSC) intercept^5^—with an intercept of one being consistent with no confounding. We estimated the intercept both before correction, and after PC loading regression correction using loadings from either four SNP PCs or 14 SNP and haploSNP PCs (see above). Out of the 36 summary statistic data sets, eight data sets, including all three height data sets, showed some evidence of confounding (intercept > 1.05). This indicates that uncorrected stratification may be a concern even in GWASes that have access to individual-level data and have applied standard methods to correct for stratification.

We first examined LDSC intercepts after PC loading regression correction using four SNP PCs (Table 3). We observed a statistically significant reduction (> 2 s.d; see Methods) in LDSC intercept^5^—from 1.35 (s.e. 0.02) to 1.24 (s.e. 0.02)—for GIANT height summary statistics, which are known to be affected by uncorrected population stratification^13^. We observed a proportionate reduction in the intercept for a previous of GWAS of human height with smaller sample size^12^. As expected, we did not observe any reduction in the intercept for 23andMe summary statistics that included SNP PC covariates.

**Table 3.** LD score intercepts in empirical summary statistic data sets before and after applying PC loading regression.

| Phenotype | Source | Uncorrected |  | SNP PCs |  | SNP + haploSNP PCs |  |
| --- | --- | --- | --- | --- | --- | --- | --- |
| Height | 23andMe | 1.41 | (0.036) | 1.40 | (0.036) | 1.16 | (0.032) |
| Height | Wood et al. 2014 Nat. Genet. | 1.35 | (0.022) | 1.24 | (0.021) | 0.99 | (0.018) |
| Height | Lango Allen et al. 2011 Nature | 1.11 | (0.016) | 1.08 | (0.015) | 1.06 | (0.015) |
| Hair grayness | 23andMe | 1.10 | (0.015) | 1.09 | (0.015) | 1.09 | (0.015) |
| Eye color | 23andMe | 1.09 | (0.025) | 1.08 | (0.024) | 0.99 | (0.023) |
| Unibrow | 23andMe | 1.06 | (0.011) | 1.06 | (0.011) | 1.06 | (0.011) |
| Motion sickness | 23andMe | 1.06 | (0.014) | 1.06 | (0.014) | 1.06 | (0.014) |
| Hair curliness | 23andMe | 1.06 | (0.015) | 1.06 | (0.015) | 1.05 | (0.015) |
| Male pattern baldness | 23andMe | 1.05 | (0.012) | 1.05 | (0.012) | 1.05 | (0.012) |
| Hair color | 23andMe | 1.05 | (0.017) | 1.04 | (0.016) | 0.97 | (0.015) |
| Ulcerative colitis | Jostins et al. 2012 Nature | 1.05 | (0.011) | 1.05 | (0.011) | 1.05 | (0.011) |
| Chin dimple | 23andMe | 1.05 | (0.011) | 1.04 | (0.011) | 1.04 | (0.011) |
| Male back hair | 23andMe | 1.04 | (0.012) | 1.04 | (0.012) | 1.03 | (0.012) |
| Crohn's disease | Jostins et al. 2012 Nature | 1.03 | (0.013) | 1.03 | (0.013) | 1.03 | (0.013) |
| Schizophrenia | PGC 2014 Nature | 1.03 | (0.014) | 1.02 | (0.014) | 1.02 | (0.014) |
| Shoe size | 23andMe | 1.02 | (0.011) | 1.02 | (0.011) | 1.02 | (0.011) |
| Skin color | 23andMe | 1.02 | (0.009) | 1.02 | (0.009) | 0.99 | (0.009) |
| Dimples | 23andMe | 1.02 | (0.010) | 1.02 | (0.010) | 1.02 | (0.010) |
| Coronary artery disease | Schunkert et al. 2011 Nat. Genet. | 1.02 | (0.010) | 1.02 | (0.010) | 1.02 | (0.010) |
| Male facial stubble | 23andMe | 1.02 | (0.010) | 1.02 | (0.010) | 1.02 | (0.010) |
| Type II Diabetes | Morris et al. 2012 Nat. Genet. | 1.02 | (0.009) | 1.01 | (0.009) | 1.01 | (0.009) |
| Male age voice deepened | 23andMe | 1.02 | (0.010) | 1.02 | (0.010) | 1.01 | (0.010) |
| Educational attainment | Rietveld et al. 2013 Science | 1.01 | (0.011) | 1.01 | (0.011) | 1.01 | (0.011) |
| Widow's peak | 23andMe | 1.01 | (0.009) | 1.01 | (0.009) | 1.01 | (0.009) |
| Nose size | 23andMe | 1.01 | (0.012) | 1.01 | (0.011) | 1.01 | (0.011) |
| Beighton hypermobility | 23andMe | 1.01 | (0.011) | 1.01 | (0.011) | 1.00 | (0.011) |
| Cup size | 23andMe | 1.01 | (0.010) | 1.01 | (0.010) | 1.01 | (0.010) |
| Female hair loss | 23andMe | 1.01 | (0.013) | 1.01 | (0.013) | 1.00 | (0.012) |
| Smoking status | TAG Consortium 2010 Nat. Genet. | 1.00 | (0.008) | 1.00 | (0.007) | 0.99 | (0.007) |
| Bipolar disorder | PGC 2011 Nat. Genet. | 1.00 | (0.012) | 1.00 | (0.012) | 1.00 | (0.012) |
| Fasting blood glucose | Manning et al. 2012 Nat. Genet. | 0.99 | (0.009) | 0.99 | (0.010) | 0.99 | (0.010) |
| Anorexia | Boraska et al. 2014 Mol. Psych.* | 0.96 | (0.010) | 0.94 | (0.010) | 0.94 | (0.010) |
| LDL | Teslovich et al. 2010 Nature* | 0.90 | (0.011) | 0.90 | (0.011) | 0.90 | (0.011) |
| Triglycerides | Teslovich et al. 2010 Nature* | 0.89 | (0.010) | 0.89 | (0.010) | 0.89 | (0.010) |
| HDL | Teslovich et al. 2010 Nature* | 0.87 | (0.011) | 0.87 | (0.011) | 0.87 | (0.011) |
| BMI | Speliotes et al. 2010 Nat. Genet. * | 0.77 | (0.009) | 0.77 | (0.009) | 0.76 | (0.009) |

\*We note that those traits with uncorrected LD score regression intercept < 1 are likely to have applied a genomic control correction.
We applied PC loading regression to 36 GWASes. Here we list the LD score regression intercepts before correction and after correction with either four SNP-PCs or 14 SNP and haploSNP PCs. Summary staistics for which we observed a nominally significant reduction LD Score intercept are listed in bold. PC loading regression produced the largest reduction for the three GWASes of human height.

We next examined LDSC intercepts after PC loading regression using 14 SNP and haploSNP PCs (Table 3). We observed a significant reduction in LDSC intercept for four 23andMe traits: height, hair color, eye color and skin color—eliminating evidence of confounding for the last three. In addition, inclusion of haploSNP PC loadings further reduced evidence of confounding in publicly available height GWAS summary statistics, yielding corrected summary statistics that showed no evidence of confounding for the largest meta-analysis of height to date^13^.

#### Analysis of corrected height summary statistics

Given that the largest reduction in LD score regression intercepts was observed for the GIANT height GWAS^13^, we investigated whether correcting for stratification using PC loading regression affected the set of genome-wide significant association results. Restricting to SNPs for which PC loadings from typed or high-quality imputed GERA SNPs were available, 386 loci contained genome-wide significant SNPs before correction (slightly smaller than the 423 loci reported in ref. 12, due to restricting to highquality PC loadings). After correction via PC loading regression using 14 SNP and haploSNP PCs, 27 loci no longer contained any genome-wide significant SNPs (see Methods). Notably, among the loci that were no longer genome-wide significant, the largest drop in association statistics occurred near the lactase (*LCT*) gene^3,25^, which is known to have been subject to strong selection pressure in Europe. Indeed, a putative target SNP for this selection, rs4988235, shows a strongly suggestive signal of association to height (*P = 6.1e-8*) that is eliminated after correction (*P=0.69*) (see Figure 1).

**Figure 1.**
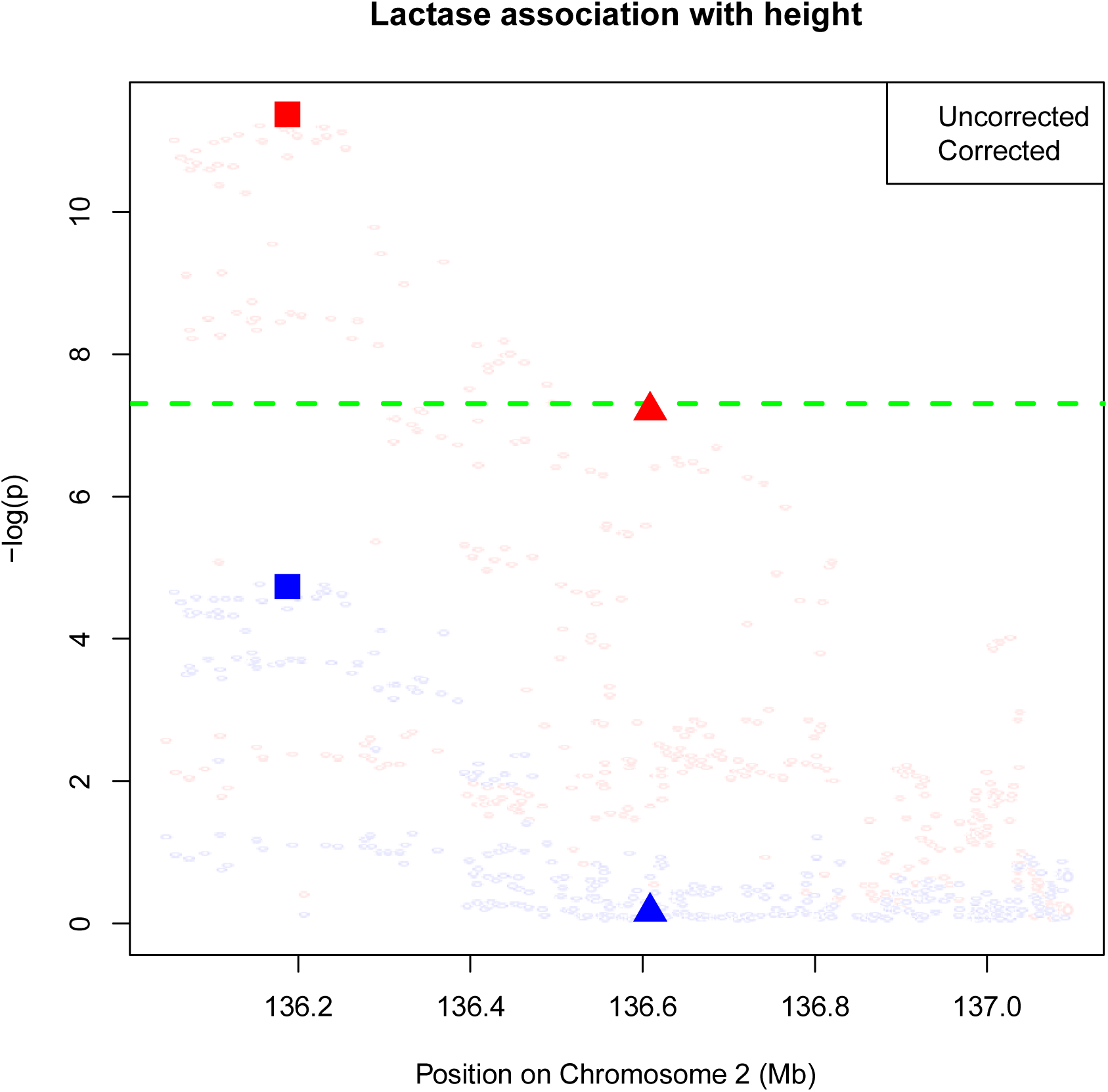
Height association at the *LCT* locus before and after PC loading regression. Here we plot the association statistics for SNPs within +/-500kb of the *LCT* gene on chromosome 2. Red and blue points are association statistics before and after correction, respectively. The bold squares indicate the most significant association prior to correction, which clearly drops below genome-wide significance. The bold triangle indicates a putatively causal SNP for lactase persistence (rs4988235), presumed to be driving allele frequency differentiation in Europe. This SNP is borderline genome-wide significant prior to correction (*P=6e-8*), and not significant after correction (*P=0.69*).

Of a total of 1.6M SNPs that were analyzed (see Table S1), 17,400 were genomewide significant (*P* < 5e-8) before correction. After correction, 6,058 of these SNPs were no longer genome-wide significant, and 99 other SNPs became genome-wide significant. This reduction in the number of genome-wide significant SNPs is expected due to both the removal of stratification and a loss in power from application of PC loading regression (see Methods). To ensure that we were performing meaningful stratification correction, we compared GIANT estimated effect sizes (before and after correction) to height summary statistics computed from 113k unrelated individuals of homogenous ancestry^26^ from the UK Biobank cohort^27^ (see Methods). We first focused on losers— SNPs that were no longer genome-wide significant after correction—and winners—SNPs that became genome-wide significant after correction. We observed that the relative uncorrected effect-sizes of losers, relative to UK Biobank, were statistically significantly larger than the relative uncorrected effect-sizes of winners (see Table S3). This is consistent with population stratification inflating uncorrected effect sizes for losers and deflating these effect-sizes for winners. By comparison, the relative corrected effect sizes of winners and losers were consistent with one another.

To further validate that our stratification correction eliminated the effects of environmental population stratification and not true genetic signal (e.g. due to polygenic selection (see Discussion)), we estimated genetic correlation^28^ using LD score regression between all of our height summary statistics and those from the UK Biobank cohort^27^ (see above). Removal of environmental population stratification from our height summary statistics would result in an increase in genetic correlation with UK Biobank summary statistics, while removal of true genetic signal would have the opposite effect. We observe a statistically significant increase in genetic correlation with improved correction (see Table S4), indicating that our correction is primarily removing the effects of environmental stratification.

## Discussion

We analyzed summary statistics from 36 GWASes (mean sample size n=88k), including 20 GWASes using 23andMe data, and observed a significantly inflated LD score regression (LDSC) intercept for several traits. This suggests that uncorrected population stratification may produce spurious inflation in summary statistics. This is of particular concern in the context of large meta-analyses in which individual-level data is not available, and standard methods to correct for stratification (i.e. inclusion of PC covariates) cannot be applied. However, uncorrected subtle stratification may also be present in analyses where SNP PC covariates were included.

Applying PC loading regression using SNP and haploSNP PC loadings estimated in the large GERA dataset^7^, we demonstrated statistically significant reductions in the LD score regression intercept^5^ for a number of traits including height, eye color, hair color and skin color—all traits that are known to be highly differentiated within Europe^29,30^. For height, these reductions were observed in both publicly available^12,13^ and 23 andMe summary statistics. Additionally, application of PC loading regression to publicly available height summary statistics eliminated a number of genome-wide significant SNPs and led to the discovery of a small set of new SNPs. Notably, the largest reduction in association statistics at eliminated loci was observed in the *LCT* region of the genome—known to be highly differentiated within Europe. For those GWAS summary statistics where the LD score regression intercept^5^ remained inflated after PC loading regression, it is likely that included PCs did not capture the relevant axes of stratification.

We note that in addition to population stratification, polygenic selection, which is thought to be acting on height in Europe^30-32^, could induce correlation between PC loadings and summary statistics. This correlation could in principle be incorrectly interpreted as population stratification and cause PC loading regression to correct away true signals. However, based on theoretical considerations we believe that any such correlation due to polygenic selection is likely to be substantially smaller than the correlation observed in our analysis of height summary statistics, and that correcting away true signals is unlikely (see Methods).

As analysis of GWAS summary statistics becomes a larger component of genetics research^28,33,34^, ensuring that these summary statistics are robust is increasingly important. Here, we provide PC loading regression as a method for producers and consumers of GWAS summary statistics to not only assess^5^, but also correct for confounding due to population stratification.

## Methods

### PC Loading Regression

Consider a phenotype *y* that is differentiated along a population axis *PC1* due to environmental stratification.

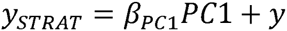

Consider a single SNP χ and its effect size, *β*, for *у* and loading *γ* for *PC1*

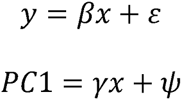

In this scenario, we can rewrite у_strat_ in terms of as χ

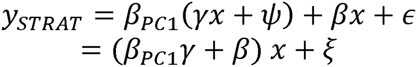

Thus, in the limit of infinite sample size the observed effect size for SNP χ on 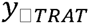 will be

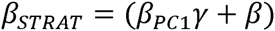

Now, let us say that we estimate summary statistics 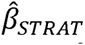 for a large number of SNPs and we are given the true SNP loadings γ. Then, if we perform a simple linear regression:

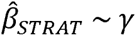

Assuming that there is no correlation between true effect sizes and SNP loadings *r(γ,β)* = 0 (see below), the estimated coefficient serves as an estimator of γ_PC1_, and the residuals of this regression:

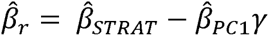

serve as estimators of the corrected coefficients *β*. We note that this can be extended to correct for an arbitrary number of principal components assuming that PC loadings are known without noise. Noise in estimates of PC loadings will limit the number of PCs that can be robustly corrected for.

#### Noisy estimated loadings 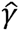 *bias estimates of* 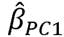

PC loadings must be estimated from a finite reference panel and statistical noise in estimates of 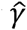 will introduce bias in the estimate of 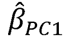

Assuming that the noise in estimates 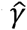 is normally distributed mean 0, and variance 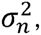, and the variance (across SNPs), in the true loadings γ is 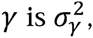, the estimated value of 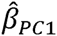 will be biased as:

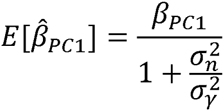

Assuming that the *PC1* perfectly separates two populations with a known *F*ST between them, 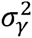 will be approximately *F*_ST_. As an example, consider structure in European Americans, where *F*_ST_ is approximately 0.005^35^. Given a sample size of *N*=50k, the expected bias will be approximately

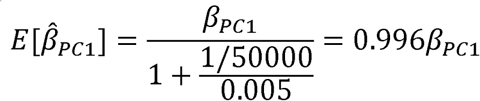

This suggests that, using available sample sizes, it will be possible to robustly estimate *β*_PC1_ and correct for population stratification of this magnitude. We note, however, that more subtle stratification (i.e. due to PCs with smaller eigenvalues) may not be adequately corrected for using loadings estimated from available reference panels. To assess our ability to correct for stratification of this magnitude we performed simulations using real PCs computed in the GERA data-set (see Results).

Application of PC loading regression can result in a loss in power due to noise introduced by the PC loadings.

#### Noisy estimated loadings 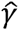 reduce power

Statistical noise in estimates of PC loadings 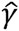 is the reduction in power that they may introduce. This loss in power will only occur if stratification exists (i.e. there is a correlation between PC loadings and estimated effect sizes) and will add noise as follows:

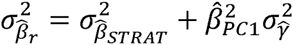

where 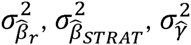 are the statistical noise in estimates of the corrected effect sizes, the uncorrected effect sizes (i.e. the original GWAS) and the PC loadings, respectively. For GWASes with small sample size—significantly smaller than the reference panel sample size (n=54k)—even a large impact of stratification will have minimal impact on study power. For GWASes with much larger sample size, as stratification becomes a larger concern the statistical noise of the PC loadings will begin to have a larger effect on the power of the corrected GWAS.

### Standardized vs. per-allele effect sizes

PC loading regression is designed for application to per-allele effect sizes from GWAS summary statistics. Under an assumption that the allele frequencies are identical in the GWAS and reference panel populations, PC loading regression can be applied to effectsizes for standardized genotypes. We test this assumption in our simulation analysis (see Table S2), and PC loading regression performed equivalently for standardized and per-allele effect sizes. Given that many GWAS summary statistics do not contain per-allele effect sizes (or sample allele frequencies so that these can be calculated), we applied PC loading regression to standardized effect sizes for all GWASes and report simulation results for standardized effect size in the Main Text (see Table 1).

### Polygenic selection on the trait of interest

If natural selection acts on the trait of interest^30,31^, it will tend to induce correlation between PC loadings and true SNP effect sizes violating the condition that *r*(*γ,β*) = 0. However, theoretically, we expect this induced correlation to be small. Consider true SNP effect sizes *β* that are correlated with SNP loadings due to polygenic selecting. Specifically, we have *β* = *c*_1_*γ* + *β*^*^, where *β*^*^ is the component of the SNP effect size that is uncorrelated with PC loadings. Now, for this phenotype, we estimate effect sizes in a GWAS with population stratification. At causal SNPs we observe estimated effect sizes:

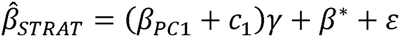

However, polygenic selection will not impact estimates at non-causal SNPs:

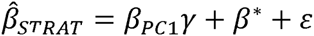

Our estimate of the contribution of environmental population stratification will be biased:

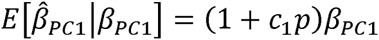

but this bias will be waited by the proportion of causal SNPs as well as the strength of selection. As such, we expect the correlation induced by natural selection to be small for realistic selection coefficients

### LD Score Regression

To assess whether the reduction of effect size variance was, in fact, due to elimination of population stratification, we applied LD Score regression^5^ using LD scores estimated from the 1000 Genomes Project^35^. Reduction in the LD score regression intercept was considered evidence that residual effect sizes were less confounded by population stratification than original effect sizes. This reduction was considered significant if it was greater than twice the standard error of the difference based on nominal standard errors. We note that this is conservative as LD score regression intercepts for corrected and uncorrected summary statistics are highly correlated and, thus, the standard error of the difference will be smaller than expected from nominal standard errors.

To assess genetic correlation between our height GWAS summary statistics and those computed from the UK Biobank, we applied LD Score regression^28^ with both constrained and unconstrained LD score regression intercepts (see Table SB2). An unconstrained LD score regression intercept should absorb the effects of stratification, reducing the improvement in estimated genetic correlation.

### Simulations Using Real Genotypes

To assess the validity of PC loading regression, we performed simulations using real genotypes in GERA data set (see below). We split the data into two partitions, each with N=27k individuals, and performed principal component analysis in each partition.

In partition 1, we simulated phenotypes stratified along PC *i* with no genetic effects.

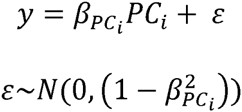

We estimated summary statistics 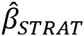 for these stratified phenotypes without including principal components as covariates.

In partition 2, we estimated the loadings 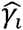 each PC *i*.

We then performed PC loading regression using summary statistics 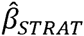, estimated in partition 1, against loadings 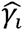 for PCs one through 10, estimated in partition 2. The resulting residuals were produced

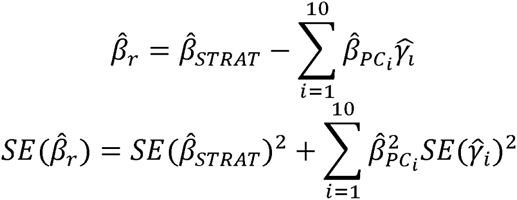

And converted to χ^2^ statistics (with 1 d.f.) as:

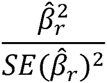

Simulations were performed including effects of stratification along each of the top four SNP PCs and each of 10 haploSNP PCs (see above). We simulated 20 phenotypes for each PC using *β*_*PC*_*i*__ = 0.1, ran LD score regression and compared the mean value of the LDSC intercept for corrected statistics and uncorrected statistics to the expected value of one under the null.

### 23andMe Dataset

For the 23andMe study, participants were drawn from the customer base of 23andMe Inc. (Mountain View, CA), a consumer genetics company^36,37^. All participants included in the analyses provided informed consent and answered surveys online according to our human subjects protocol, which was reviewed and approved by Ethical & Independent Review Services, a private institutional review board. Samples were genotyped on one of four genotyping platforms. The V1 and V2 platforms were variants of the Illumina HumanHap550+ BeadChip, including about 25,000 custom SNPs selected by 23andMe, with a total of about 560,000 SNPs. The V3 platform was based on the Illumina OmniExpress+ BeadChip, with custom content to improve the overlap with our V2 array, with a total of about 950,000 SNPs. The V4 platform in current use is a fully custom array, including a lower redundancy subset of V2 and V3 SNPs with additional coverage of lower-frequency coding variation, and about 570,000 SNPs.

Participants were restricted to a set of individuals who have >97% European ancestry, as determined through an analysis of local ancestry^38^. A maximal set of unrelated individuals was chosen for each analysis using a segmental identity-by-descent (IBD) estimation algorithm^39^. Individuals were defined as related if they shared more than 700 cM IBD, including regions where the two individuals share either one or both genomic segments identical-by-descent. This level of relatedness (roughly 20% of the genome) corresponds approximately to the minimal expected sharing between first cousins in an outbred population.

Participant genotype data were imputed against the March 2012 “v3” release of 1000 Genomes reference haplotypes, phased with ShapeIt2^40^. Data were phased and imputed for each genotyping platform separately. Data were phased using a 23andMe developed phasing tool, Finch, which implements the Beagle haplotype graph-based phasing algorithm^41^, modified to separate the haplotype graph construction and phasing steps.

In preparation for imputation, phased chromosomes were split into segments of no more than 10,000 genotyped SNPs, with overlaps of 200 SNPs. SNPs with Hardy-Weinberg equilibrium P<10−^20^, call rate < 95%, or with large allele frequency discrepancies compared to European 1000 Genomes reference data were excluded. Frequency discrepancies were identified by computing a 2x2 table of allele counts for European 1000 Genomes samples and 2000 randomly sampled 23andMe customers with European ancestry, and identifying SNPs with a chi squared P<10−15. Each phased segment was imputed against all-ethnicity 1000 Genomes haplotypes (excluding monomorphic and singleton sites) using Minimac2^42^, using 5 rounds and 200 states for parameter estimation.

The genetic association tests were performed using either linear or logistic regression as required assuming an additive model for allelic effects and controlled for age, sex, and five principal components of genetic ancestry.

### GERA Dataset

The GERA dataset includes 62,318 individuals from Northern California typed on a European-specific 670,176 SNP array^7^. To perform PCA, we focused on a previously described subset of 54,734 unrelated individuals of European ancestry^6^. We chose this data set as our reference panel for computation of PC loadings based on its large sample size and through representation of individuals of diverse European ancestry. PCA was performed on an LD pruned set of 162,335 SNPs using the fast principal component analysis feature of EIGENSOFT^4,6^ and loadings for the top four PCs were estimated on the full set of 608,981 post-QC SNPs. Separately, we computed loadings for haploSNP CRM PCs that were computed from rare haploSNPs in three bins: [0.0001-0.001),[0.001-0.005),[0.005-0.01). To assess the ability of PC loading regression to correct for population stratification, we split the full set of individuals into two halves, each containing 27,367 individuals and performed PCA (on SNPs and haploSNPs) in each half independently. We estimated summary statistics for simulated, stratified phenotypes in the first half, and used loadings computed from the second half to correct for population stratification.

### GERA Imputation

To expand the set of SNPs for which PC loadings were available, we performed imputation using the University of Michigan imputation server, using Eagle to phase genotypes and used the Haplotype Reference Consortium (r1.1) as the imputation reference panel. Because the imputation server could process only data-sets of <15k individuals, we split the data-set into four parts. We restricted to high quality imputed 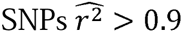 in all four parts, and computed loadings at 7.0M high-quality imputed SNPs.

To assess the quality of the PC loadings at high 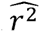 imputed SNPs, we masked 10% (or 61k) of the genotyped SNPs and only uploaded 547k SNPs to the imputation server. After imputation, we compared PC loadings at masked imputed markers to the loadings at the corresponding genotyped markers. We did this for all SNP and haploSNP PCs. We observed high correlations (> 0.9) for all PCs except for 2 haploSNP PCs (see Table 2). These PCs were excluded from subsequent analysis.

### Analysis of genome-wide significant loci

We analyzed each of the 423 loci published in Wood et al. (ref. 12), for marginal association after merging with well-imputed GERA SNPs. We analyzed all SNPs that were within 500kb of each locus. If no SNPs were found to be genome-wide significant the locus was considered not genome-wide significant. This was repeated for summary statistics before and after correction using PC loadings. For the purpose of association analysis, we used original GIANT height summary statistics. For all other analyses, we used re-inflated height summary statistics to undo the effects of genomic control^43^ as previously described^5^.

### UK Biobank height association analysis

As a validation of our correction for stratification, we compared corrected and uncorrected height summary statistics to height summary statistics computed from 113k unrelated individuals of homogenous ancestry^26^ from the UK Biobank cohort^27^. Summary statistics were estimated for 20.0M imputed SNPs after filtering SNPs that had MAF < 0.1%. Statistics were estimated using standard linear regression implemented in the BOLT-LMM^44^ software, including principal components computed on these same individuals using an LD pruned subset of SNPs^26^.

## Acknowledgements

We would like to thank the research participants and employees of 23andMe, particularly David Hinds and Adam Auton, for making this work possible. This research was funded by NIH grants R01 HG006399, R01 MH101244 and U01 HG009088. This research has been conducted using the UK Biobank Resource.

